# A high-throughput assay quantifies thermal scaling of *Drosophila* development with minute-scale precision

**DOI:** 10.64898/2026.03.06.710149

**Authors:** Daniel Sobrido-Cameán, Fernando Claro-Linares, Noelia Ruiz-Gómez, Patricia Rojas-Ríos, María Olmedo

## Abstract

Precise regulation of developmental timing is essential for coordinated growth and robust development, yet staging remains technically challenging in many model systems. In *Drosophila melanogaster*, developmental timing has traditionally been assessed using low-throughput or coarse staging methods, limiting insight into how individual larval stages respond to environmental and genetic perturbations. Here, we present a high-throughput, real-time luminometry assay that enables continuous, automated measurement of postembryonic development in individual *Drosophila* larvae. By monitoring feeding-dependent luciferase activity, this method reliably detects transitions between larval instars and molts of individual larvae with minute-scale temporal resolution. Using this platform, we provide a quantitative, minute-precision, stage-resolved description of larval development across large cohorts, revealing distinct patterns of variability and weak temporal coupling between stages. We next examine how temperature shapes developmental timing across a broad thermal range. Increasing temperature uniformly accelerates larval development while preserving the proportional contribution of each stage, indicating that the temporal architecture of development is maintained as overall pace changes. Developmental rate follows predictable thermodynamic scaling within a defined temperature window, enabling precise estimation of thermal parameters at both whole-organism and stage-specific levels. Finally, we demonstrate that the assay is compatible with genetic perturbations. This work establishes a scalable, high-precision framework for measuring postembryonic development in *Drosophila melanogaster*. By revealing the modular and coordinated nature of larval growth, it enables systematic dissection of genetic, metabolic, and environmental control of developmental timing and provides a platform to explore fundamental principles of development and robustness across species.

## Introduction

Precise quantification of developmental timing is fundamental for understanding how genetic, cellular, and environmental factors shape organismal growth. Development unfolds through a series of discrete transitions whose timing is tightly regulated across metazoans. The duration of developmental stages reflects the integration of intrinsic genetic programs with extrinsic cues, and even modest shifts in tempo can reflect changes in environmental conditions, energy availability, or endocrine state (Busby et al., 2024; Ferree et al., 2016; Garcia-Ojalvo & Bulut-Karslioglu, 2023; Iwata & Vanderhaeghen, 2024). Developmental timing is thus not merely a passive clock but a dynamic system shaped by interactions between gene regulatory networks, hormonal pulses, and metabolic constraints (Shingleton & Frankino, 2018; Tennessen & Thummel, 2011). In insects, developmental progression is organized around a sequence of molts and instars gated by the endocrine system. Pulses of ecdysone coordinate transitions in growth, linking environmental inputs such as nutrition and temperature to the activation of stage-specific gene expression programs Because most insects are poikilotherms, temperature directly governs developmental pace at the levels of growth, endocrine signaling, and metabolism. This tight coupling makes holometabolous insects an ideal system for understanding how environmental factors constrain developmental tempo (Chong et al., 2018; Kuntz & Eisen, 2014; Maulana et al., 2022; Sekajova et al., 2022; Zuo et al., 2012).

*Drosophila melanogaster* is one of the most powerful and widely used model organisms in developmental biology. Its rapid life cycle, genetic tractability, and deeply characterized developmental stages have made it essential for uncovering fundamental principles of pattern formation, growth control, organogenesis, and environmental modulation of developmental programs (Giansanti et al., 2025; Roberts, 2006; Tolwinski, 2017). In *D. melanogaster*, developmental timing has been studied for nearly a century, yet most classical analyses measured only total developmental duration at broad temperature intervals (Al-Saffar et al., 1996; Powsner, 1935; Robinson & Partridge, 2001). These foundational studies established that development accelerates with temperature, but provided limited resolution on how individual larval stages contribute to whole-organism thermal responses. Moreover, whether all stages scale uniformly with temperature, and whether specific transitions deviate from canonical thermodynamic expectations, has remained unclear. Although developmental rates accelerate with temperature in ectotherms, the mechanisms that coordinate this acceleration across tissues and developmental modules remain poorly understood. Emerging evidence suggests that distinct developmental stages may differ in their sensitivities to environmental factors (Mata-Cabana et al., 2022), raising the possibility that developmental tempo is governed by multiple, stage-specific processes rather than a universal timer. High-resolution quantification of developmental timing across development is therefore essential to understand how organisms maintain coordinated development under changing environmental conditions. Although some visual tracking approaches exist (Schumann & Triphan, 2020), *Drosophila* studies of developmental timing still rely largely on manual or low-throughput methods. Continuous, high-temporal-resolution measurement of individual larvae has been lacking, making subtle shifts due to environmental, genetic, or metabolic perturbations difficult to detect. Our high-throughput luminometry assay addresses this gap, providing real-time, quantitative monitoring of developmental progression across large cohorts.

In *Caenorhabditis elegans*, we addressed this limitation by developing a luminescence-based, high-throughput method that continuously monitors developmental progression in individual animals (Olmedo et al., 2015). This approach leverages feeding-dependent luciferase activity to detect stage transitions automatically, enabling large-scale, precise measurements of larval development (Mata-Cabana et al., 2020, 2022; Olmedo et al., 2015, 2020).To validate the method and illustrate its utility in *Drosophila*, we focused on a classical question in biology: how temperature controls developmental speed. Several studies propose that developmental rates across species follow predictable thermal scaling rules, like the Arrhenius equation (Boukal et al., 2015; Gillooly et al., 2002; Jarośík et al., 2004; Zuo et al., 2012), yet the extent to which distinct developmental stages share similar temperature sensitivities remains unresolved. In *Drosophila*, embryogenesis has been reported to scale uniformly with temperature (Kuntz & Eisen, 2014), suggesting a universal biochemical rate-limiting step. Yet, despite precise measurements in embryogenesis, larval stages remain largely unexplored, limiting our understanding of stage-specific thermal sensitivities.

Here, we introduce a high-throughput, real-time assay to quantify postembryonic development in *D. melanogaster*. This method adapts the principles of the *C. elegans* luminescence assay to the fruit-fly, enabling automated staging of large cohorts with minute-scale resolution. By precisely defining the temperature window in which development follows the Arrhenius equation, estimating lower developmental thresholds (LDT) and sums of effective temperatures (SET) at the level of single larvae, we provide a framework for dissecting the biochemical, genetic, metabolic, and endocrine mechanisms that coordinate growth and development. Moreover, because both nematodes and insects are ecdysozoans that share molting-based developmental transitions, this framework offers the potential to extend automated developmental timing assays to other species within the clade. Together, this work establishes a platform for quantitative, stage-resolved measurement of larval development in *Drosophila*, enabling systematic studies of environmental modulation, genetic perturbations, and cell-intrinsic regulatory mechanisms.

## Results

### High throughput, automated analysis of *Drosophila melanogaster* developmental timing

With the aim of quantifying developmental timing in *Drosophila*, we have adapted a method originally developed for *C. elegans* (Olmedo et al., 2015). This method is based on the use of larvae that express the enzyme luciferase from *Photinus pyralis*. Luciferase catalyzes the oxidation of luciferin, by a reaction that emits light. In the assay, luciferin is provided with the food, so that is only available during intermolts, when larvae are feeding. During molting periods, the larval mouth is blocked by a cuticular plug and luciferin intake ceases. This way, larvae emit luminescence only during intermolts, allowing the identification of the transitions between intermolts (I) and molts (M) over the four stages of *C. elegans* larval development (Olmedo et al., 2015). To test the method in *D. melanogaster*, we used flies ubiquitously expressing luciferase from *Photinus pyralis* and *Renilla reniformis*. We placed one age-matched embryo (0 to 3 hours old) per well of a 96-well plate and recorded luminescence (Figure 1a). The signal increased at the moment of hatching and stayed high for most of larval development, except for two interruptions coinciding with the first and second molts (Figure 1a, b). After the third instar, the signal stays very low during the pupal stage, which does not feed. This result demonstrates that the luminescence-based assay can reliably track developmental transitions in *Drosophila* larvae, allowing precise identification of molts and intermolts throughout postembryonic development.

**Figure 1.**
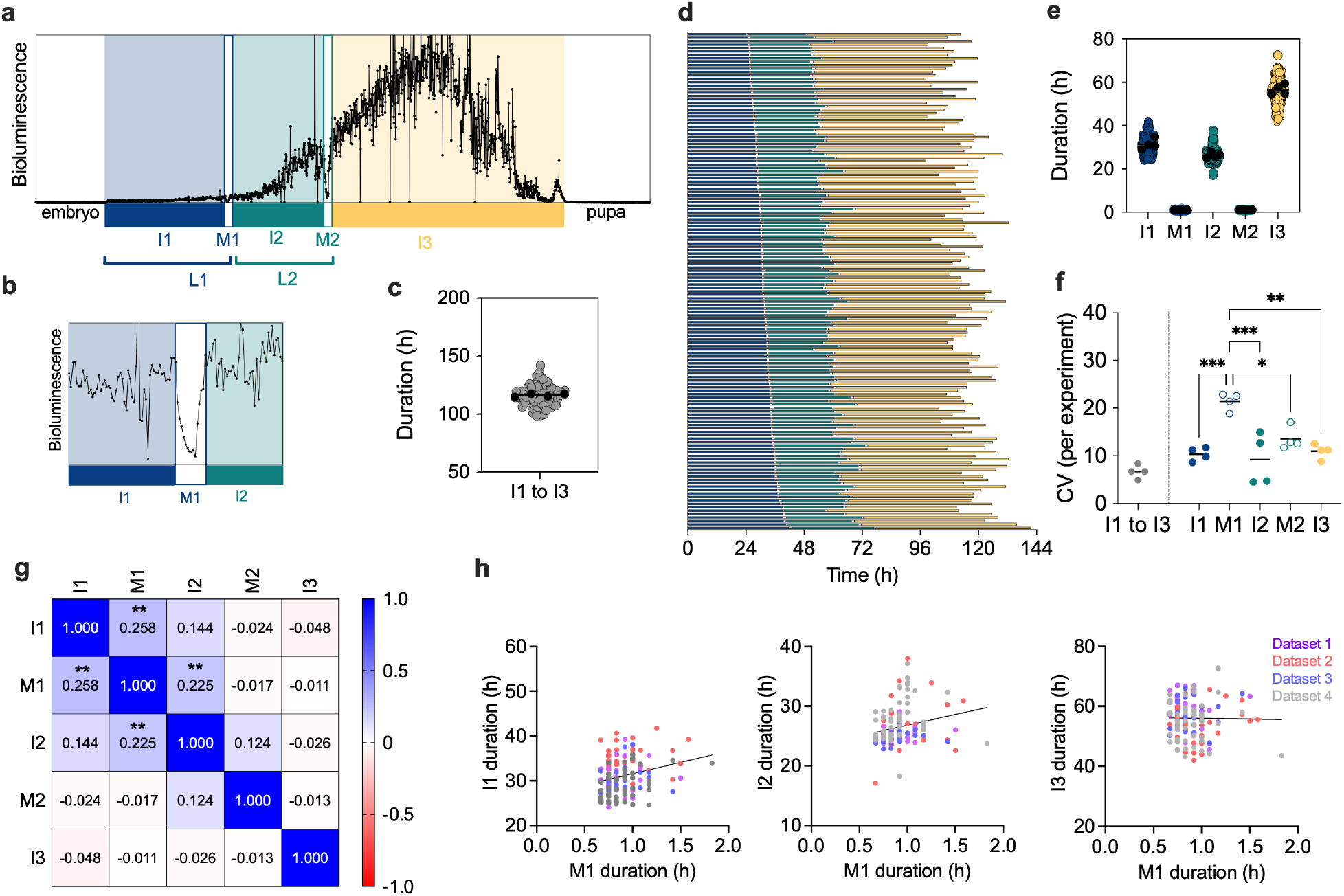
Luminescence signal reports larval development. (a) Representative luminometry trace from a single larva expressing luciferase, illustrating continuous bioluminescence during feeding periods and sharp signal reductions during molts. (b) Magnified view of the first molt (M1), showing the characteristic transient drop in luminescence that marks cessation of feeding and enables precise identification of molt transitions. (c) Distribution of total postembryonic developmental time (hatching to pupariation) across all larvae at standard temperature (25 ºC). The mean of each dataset is shown as black dots. (d) Individual developmental trajectories of age-matched larvae reared at 25 °C, showing the duration of each larval stage (I1–I3) and associated molts (M1–M2) for 145 animals measured in 4 independent experiments. (e) Quantification of the duration of each intermolt and molt at 25 °C, showing with black dots the mean stage duration for each dataset. (f) Coefficient of variation (CV) for total postembryonic development and for each larval stage, calculated across four independent experiments, illustrating differences in temporal precision among stages. (g) Pairwise correlation matrix for all combinations of developmental stages showing the Pearsons value and including p values when significant. ** p < 0.01. (h) Pairwise correlations between the durations of M1 vs all instars.

### Quantitative analysis of *Drosophila melanogaster* postembryonic development

We measured the duration of all stages from hatching to pupariation for 145 larvae raised at 25 ºC, the standard rearing temperature for *D. melanogaster* (Figure 1 c-e). The average (± SD) duration of development from hatching to pupariation in four independent experiments was 115.50 (± 8.27) hours (Figure 1c). The duration of each stage was 31.11 (± 3.93) hours for intermolt 1(I1), 0.92 (± 0.20) hours for M1, 26.49 (± 3.20) hours for I2, 1.04 (± 0.15) hours for M2 and 55.97 (± 6.51) hours for I3 (Figure 1d, e).

To assess the precision of developmental timing, we calculated the coefficient of variation (CV) for each stage and for the total duration in each of the four independent experiments (Figure 1f). We observed that temporal variability differs between postembryonic developmental stages. In particular, molting phases displayed lower precision compared to intermolt periods, with M1 showing the highest CV, indicating greater inter-individual variability during this transition. In contrast, the intermolts (I1-I3) showed the lowest CV, indicating high precision during these developmental stages. These results suggest that specific developmental transitions are more tightly regulated than others, potentially revealing the presence of robust timing control mechanisms during molting events.

We further calculated the fraction of development represented by each larval stage by dividing the duration of each stage by the total postembryonic developmental time. This analysis provides insight into how developmental time is allocated across stages, revealing whether certain instars or molts disproportionately contribute to overall development. The mean fractions across all larvae were 0.2696 (± 0.0296) for I1, 0.0095 (± 0.0016) for M1, 0.2296 (± 0.0253) for I2, 0.0096 (± 0.0011) for M2 and 0.4840 (± 0.0380) for I3 (Supplementary Figure 1a). These results indicate that the I3 constitutes the largest proportion of postembryonic development, while the molt stages (M1 and M2) represent only a minimal fraction of the total developmental time. Finally, to investigate potential correlations between the durations of different larval stages within individual animals, we performed pairwise comparisons (Figure 1g, h; Supplementary Figure 1b, c). These analyses revealed modest but detectable correlations between the duration of M1 with I1 and I2.

Together, these results demonstrate that luminometry provides a robust, high-resolution readout of postembryonic development in *Drosophila*. The method allows precise quantification of stage durations, assessment of inter-individual variability, and analysis of developmental coupling across larval instars, opening the door for high-throughput studies of genetic and environmental modulators of developmental timing.

### Temperature dependence of larval development

Having validated that our method provides precise, stage-resolved developmental time, we next applied it to examine how temperature shapes larval development. Although the inverse relationship between temperature and developmental time in *D. melanogaster* is well established, previous work has largely relied on coarse staging (Lee et al., 2019; Turingan et al., 2024). By revisiting this question with high temporal resolution and detailed segmentation of all instars and molts, we aimed not simply to confirm established trends, but to uncover whether temperature scales all stages uniformly, alters stage proportionality, or reveals hidden structure in how individual developmental processes respond to thermal variation. We first quantified the absolute duration of each larval stage across eight temperatures (ranging from 16 to 30 °C). As expected, developmental time decreased with increasing temperature up to 28.5 ºC and increased again at higher temperature. Additionally, our high-resolution data revealed how each stage compresses across the thermal gradient (Figure 2a). These measurements provide the essential baseline: if temperature accelerates some stages disproportionately, this would imply distinct rate-limiting physiological processes. Establishing this baseline allowed us to test whether developmental acceleration is globally coordinated or stage-specific. To address this, we next calculated the fraction of total development represented by each stage at each temperature (Supplementary Figure 2a). This analysis is crucial because absolute time can mask proportional changes, an organism may develop faster overall while still redistributing time across stages. We observed that the fractional contribution of each instar remained nearly constant across temperatures, indicating that the temporal architecture of larval development is preserved even as the entire timeline expands or contracts (Supplementary Figure 2a-d). This suggests that larval development scales almost isometrically with temperature, implying strong internal coordination among stages.

**Figure 2.**
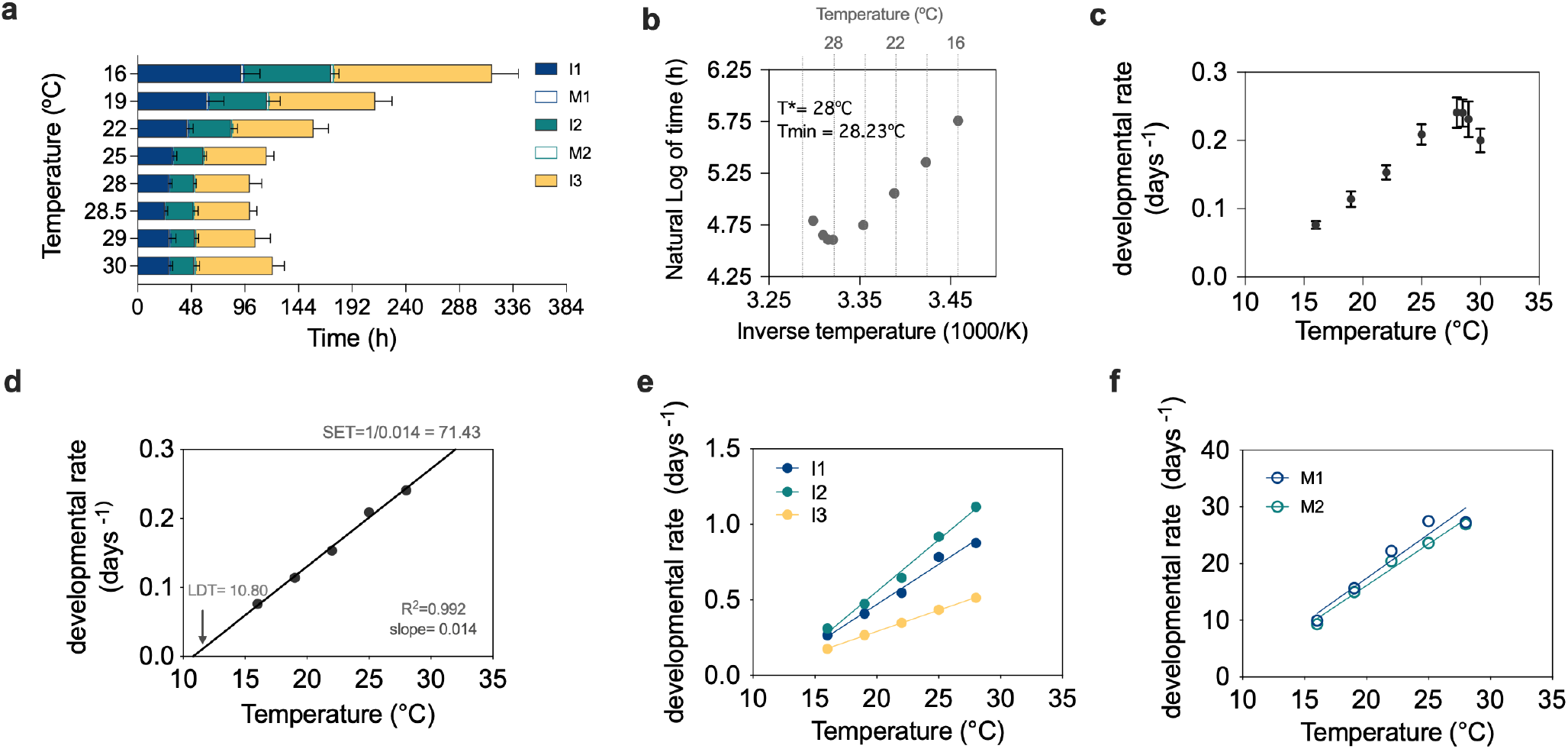
Temperature dependence of larval development in *Drosophila melanogaster*. (a) Mean duration of each developmental stage across temperatures illustrating the distribution of developmental tempo across thermal conditions. (b) Arrhenius plot (natural logarithm of time vs inverse temperature in kelvin) for the duration of development (I1 to I3). (c) Relationship between overall developmental rate (days−1) and temperature, showing the acceleration of development at higher temperatures. (d) Plot of developmental rate using only temperatures within the linear range (16 ºC-28 ºC); linear regression was used to estimate the lower developmental threshold (LDT) and the sum of effective temperatures (SET). (e) Plot of developmental rate for individual larval instars within the temperatures that follow the Arrhenius relationship. (f) As in panel e, but for molt transitions.

Next, we examined whether the overall pace of postembryonic development can be described by a biophysical model. In order to perform the corresponding analysis for postembryonic development, we plotted the natural logarithm of the duration against the inverse of the temperature in Kelvin (K) (Figure 2b). This representation allowed us to test whether larval development follows an Arrhenius-type relationship, a hallmark of processes driven by biochemical reaction kinetics (Arrhenius, 1889). Indeed, the linear correlation we observed is consistent with classical biophysical expectations. However, developmental processes typically conform to the Arrhenius equation only within a bounded thermal range, beyond which enzymatic and physiological systems begin to fail. To identify the upper temperature range over which *Drosophila* larval development conforms to Arrhenius behaviour, we applied a breakpoint analysis based on deviations from linear Arrhenius scaling. This approach allowed us to estimate the temperature limit at which the Arrhenius approximation remains valid (T*) and the temperature that sustains the fastest development (Tmin). We found that developmental rate followed Arrhenius scaling up to 28°C, whereas at higher temperatures the relationship deviated from linearity, indicating the onset of thermal stress–induced physiological constraints. The fastest development occurs at 28.23ºC (Figure 2b).

Next, we analysed the variation of developmental rate by plotting the developmental rate (days^−1^) against temperature (Fig 2c). The increase of developmental rate with temperature is linear up to 28°C, which coincides with the T*. Therefore, all subsequent analyses of developmental rate were restricted to the temperature range below 28 ºC. Within this range, linear regression of the developmental rate against temperature allows estimates of the lower developmental threshold (LDT) and the sum of effective temperatures (SET; Figure 2d). Because whole-organism developmental rate reflects the integration of multiple underlying processes, we examined whether the developmental rate of individual stages also follows a linear correlation with temperature. Both instars and molts follow a linear Arrhenius relationship over the permissive temperature range (Figure 2e, f). These analyses reveal that both growth phases (instars) and transitional events (molts) obey predictable thermodynamic scaling, reinforcing that the entire larval program responds to temperature in a coordinated and energetically constrained manner.

Together, these findings elevate a classical observation, temperature modulates developmental speed, into a detailed quantitative framework. By dissecting each component of the larval program across a broad thermal range, we show that temperature uniformly rescales development while preserving its internal structure, offering a foundational dataset for probing how metabolic and environmental signals jointly shape developmental timing.

### Developmental timing under genetic perturbation

Building on the precision of our automated developmental measurements, we next applied the method to genetically manipulated animals, taking advantage of *D. melanogaster’s* powerful genetic toolkit. To this end, we reproduced a well-characterized genetic manipulation known to affect developmental timing by reducing TOR signaling through ubiquitous co-expression of the TSC1 and TSC2 inhibitory complex using the UAS-TSC1/2 line, as described by Layalle et al. (2008) (Layalle et al., 2008). In that study, TOR downregulation delayed the third larval instar without substantially affecting earlier stages. For this purpose, we misexpressed TSC1/2 ubiquitously using the *daughterless*-GAL4 driver and combined it with our *ubiquitous* Luciferase-expressing background, allowing us to record developmental luminescence across all larval stages using our high-throughput assay

(Figure 3a). Our results recapitulated the published phenotype: the third intermolt was specifically extended, while earlier stages were largely unchanged (Figure 3b-f). The average duration of I3 was 52.38 (± 7.50) hours for control (daughterless-GAL4, ubiquitous-Luciferase/+) and 72.95 (± 14.81) hours for UAS-TSC1/2 (daughterless-GAL4, ubiquitous-Luciferase/UAS-TSC1/2). Importantly, our method enabled rapid, automated, and quantitative analysis across dozens of individuals in parallel, confirming the developmental delay with high temporal precision that also resolved the duration of the molts. These findings demonstrate that our assay is fully compatible with genetically modified lines, providing a fast, reliable, and precise tool to probe the effects of specific genetic perturbations on developmental timing. This approach therefore has broad applicability for the *Drosophila* research community, facilitating high-throughput investigations of genetic and environmental modulators of development.

**Figure 3.**
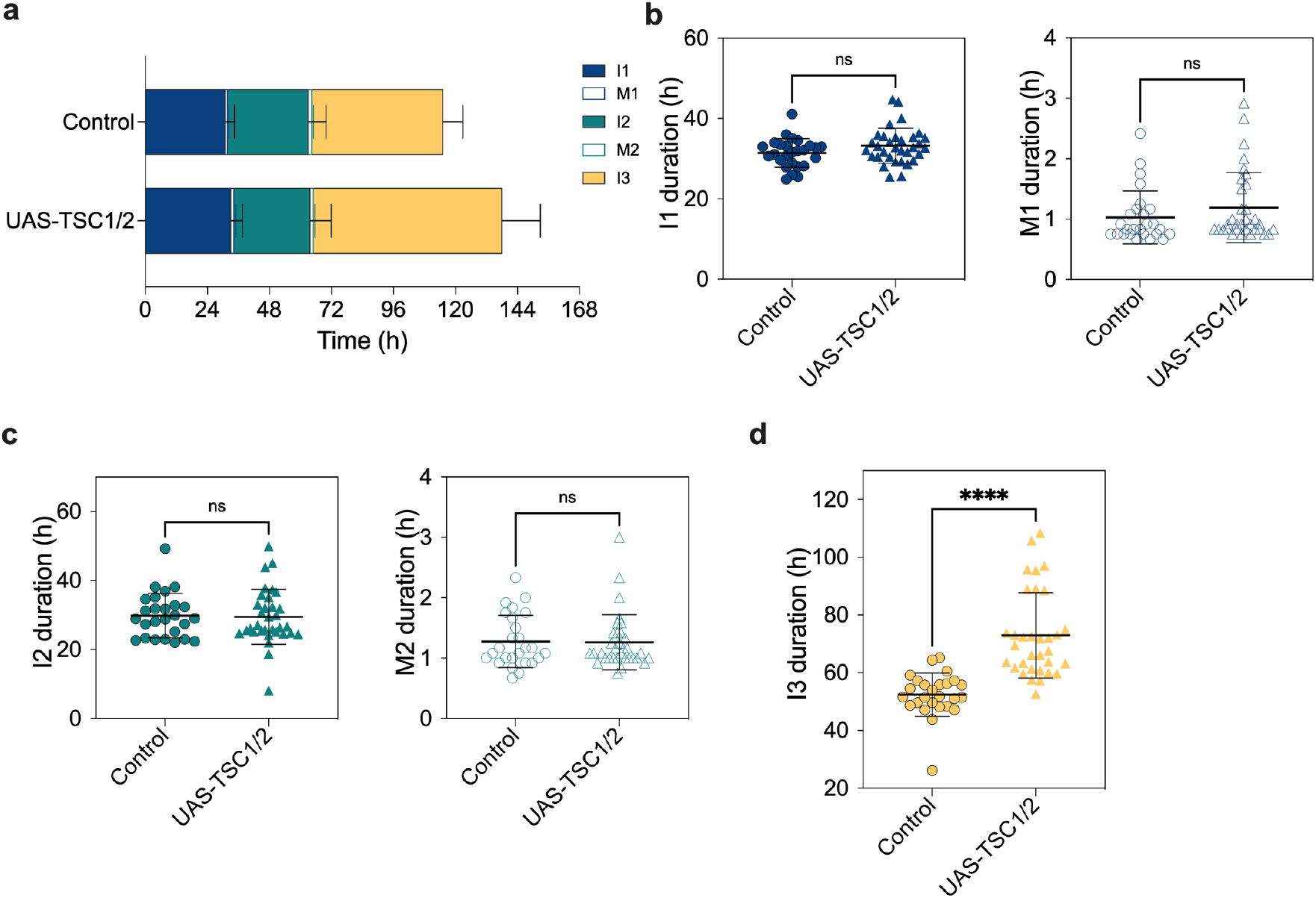
Ubiquitous TSC1/2 misexpression selectively delays the third larval instar. (a) Mean duration of each developmental stage across temperatures illustrating the distribution of developmental tempo for the control group (*daughterless*-GAL4, Luciferase/+) vs the UAS-TSC1/2 (*daughterless*-GAL4, Luciferase/UAS-TSC1/2). (b) Quantification of the duration of I1 and M1 of control group vs UAS-TSC1/2. (c) Quantification of the duration of I2 and M2 of control group vs UAS-TSC1/2. (d) Quantification of the duration of I3 of control group vs UAS-TSC1/2 showing significant differences, p-value < 0.0001.

## Discussion

How animals regulate developmental timing is a fundamental question in developmental biology, with broad implications for evolutionary ecology and the systems-level coordination of growth and metabolism. Although timing of *Drosophila* embryogenesis has been quantified with remarkable precision (Kuntz & Eisen, 2014), the postembryonic period, which constitutes the majority of development time, has lacked a high-resolution assay. Here, we adapted a luciferase-based luminometry approach originally developed for *C. elegans* (Olmedo et al., 2015) to quantitatively track *D. melanogaster* larval development at high temporal resolution. Our results validate this method in an insect model for the first time and reveal new biological insights into the structure, variability, and thermal scaling of the larval trajectory. Both *C. elegans* and *D. melanogaster* belong to the superphylum Ecdysozoa, whose members grow through ecdysis, i.e., molting of the cuticle. This suggests that the same method could likely be applied to other ecdysozoans, such as tardigrades, for which genetic manipulation tools are beginning to be developed (Kondo et al., 2024).

### A scalable, stage-resolved method to quantify *Drosophila* postembryonic development

By following luciferase activity in individual animals across development, we show that the luminometric signal reliably marks transitions between intermolts and molts, enabling automated identification of all three larval instars and the two molts (Figure 1a). The method is reproducible across experiments (Figure 1), and implemented in high-throughput formats (96 wells plate), offering unprecedented scalability for genetic or pharmacological screens targeting developmental timing. Previous approaches relied on invasive sampling, low temporal resolution, or labor-intensive manual staging (Chen et al., 1996; Juarez-Carreño & Geissmann, 2023; Layalle et al., 2008; Powsner, 1935). Our approach provides continuous, quantitative data from single larvae with a high degree of automation that fundamentally improves the accuracy, throughput, and reproducibility of postembryonic staging. This technological advance enables new types of analyses that were previously infeasible, such as quantifying the variance structure of stage durations, assessing correlations between stages within individuals, and measuring thermodynamic properties of developmental rate with fine resolution.

### Stage-specific variability reveals modular control of developmental tempo

A key finding is that temporal precision varies significantly between stages. First and second instars display relatively low variability, whereas molts, particularly M1 is more heterogeneous across individuals (Figure 1f). This suggests that different stages are governed by regulatory modules with distinct sensitivity to internal and external factors. Intermolt stages are dominated by continuous feeding, exponential growth, and biosynthetic demand, processes that depend on reaching specific mass, nutritional, and metabolic thresholds before progression is permitted (Tennessen et al., 2014; Tennessen & Thummel, 2011). These thresholds impose relatively fixed requirements for cell growth, nutrient assimilation, and metabolic flux, thereby constraining the extent to which stage duration can vary between individuals. By contrast, molts show greater inter-individual variability because they are non-feeding, hormonally driven transitions whose timing depends on endocrine checkpoints rather than metabolic thresholds. Molting requires precise coordination of ecdysone pulses, cuticle synthesis, apolysis, and behavioural programs for ecdysis (Mirth & Riddiford, 2007; Yamanaka et al., 2013). Variability in molt duration arises from differences in ecdysone amplitude and timing, tissue sensitivity to hormonal cues, and stochastic fluctuations in cuticle synthesis and epidermal remodeling (King-Jones et al., 2005; Morrow & Mirth, 2024; Rewitz et al., 2013; Shimell et al., 2018; Yamanaka et al., 2013). Together, these observations reinforce the idea that postembryonic development comprises phases governed by distinct regulatory mechanisms.

### Larvae differ in global tempo, but stages are not fully coupled

By examining correlations between the durations of different instars, we find modest but detectable coupling, particularly across earlier stages (Figure 1g). However, these correlations are weak and indicate that variation in development time of one stage does not correlate with duration of other stages (Figure 1g). This supports models of partially independent stage regulation, in which endocrine signals (e.g., ecdysone pulses) and metabolic checkpoints can buffer stochastic fluctuations (Morrow & Mirth, 2024; Rewitz et al., 2013). Weak temporal coupling between stages may also allow larvae to adjust development dynamically in response to environmental shifts, contributing to phenotypic robustness.

### Temperature accelerates development but preserves proportional scaling

Although the overall effect of temperature on *Drosophila* developmental time is well known (Chong et al., 2018; Powsner, 1935), our single-animal, stage-resolved dataset provides unprecedented precision. Across the 16 to 28 °C range, all instars shorten markedly with increasing temperature, yet the fraction of total development occupied by each stage remains constant (Figure 2a; Supplementary Figure 2a-c). This proportional scaling echoes classical observations that embryonic processes scale uniformly with temperature (Kuntz & Eisen, 2014) and now extends this principle to the entire larval period. Such proportionality suggests that different developmental modules share upstream rate-limiting constraints, potentially metabolic (Tennessen et al., 2014), thermodynamic (Jarośík et al., 2004), or endocrine (Mirth & Riddiford, 2007). The consistency of stage fractions across temperatures reinforces the idea that development progresses through a coordinated temporal framework.

### Determination of the Arrhenius-consistent thermal range

A key advance of this work is the precise quantification of the temperature range in which *Drosophila* larval development adheres to the Arrhenius equation (Figure 2c). Developmental rate increased exponentially with temperature at moderate ranges, but deviated sharply at higher temperatures (Figure 2c). To identify the threshold beyond which Arrhenius no longer applies, we applied a breakpoint analysis, revealing that temperatures above 28 °C induce systematic deviations. Breakdown of Arrhenius scaling at high temperatures likely reflects physiological limits, such as protein misfolding, impaired feeding, or endocrine disruption. By contrast, the lower-to-mid temperature range exhibits clean thermodynamic scaling, enabling accurate estimation of the LDT and the SET, parameters widely used in insect developmental ecology (Boukal et al., 2015; Jarośík et al., 2004; Jarošík et al., 2002; Mata-Cabana et al., 2022).

### High-throughput genetic analysis of developmental timing

Our ability to recapitulate a well-established, stage-specific developmental delay caused by ubiquitously co-expression of TSC1/2 further demonstrates the general applicability of this method to genetically manipulated animals (Figure 3). Unlike manual staging approaches, our assay captures developmental timing continuously, automatically, and at high temporal resolution, enabling precise quantification of how specific genetic perturbations redistribute time across larval stages. Importantly, the selective extension of the third instar was readily resolved without prior assumptions about which stages were affected, highlighting the power of unbiased, whole-development measurements. Given the extensive genetic toolkit available in *Drosophila*, this approach provides a scalable platform for systematic genetic screening and dissection of genetic, metabolic, and environmental regulators of developmental timing.

Together, this work establishes developmental timing in *D. melanogaster* as a quantitatively accessible, genetically tractable trait that can be measured with high temporal resolution across entire postembryonic development. By combining continuous luminometric readouts with stage-resolved analysis, we reveal that larval development is organized as a coordinated yet modular program, whose overall pace can be uniformly rescaled by temperature while remaining sensitive to stage-specific genetic perturbations. In doing so, we establish a new standard for measuring developmental time in *Drosophila*, enabling long-standing questions about developmental tempo, stage coupling, metabolic control, and thermal scaling to be addressed with unprecedented precision. This framework bridges classical physiological models of development with modern genetic and systems-level approaches, providing a foundation for future studies of growth control, plasticity, and robustness in developing insects.

## Materials and Methods

### *Drosophila* maintenance and stocks

*Drosophila* stocks were kept at 25 °C on standard medium. For all experiments where crosses were necessary, stocks were selected and combined as a 1:2 ratio males to females. The following fly strains were used: *ubiquitous-Luciferase* (Bloomington *Drosophila* Stock Center (BDRC) *#84127; RRID:BDSC_84127*), *daughterless*-GAL4 (BDRC #55851; RRID:BDSC_55851), *w1118* (BDRC # 3605; RRID:BDSC_3605) and *UAS-TSC1*; *UAS-TSC2* (*UAS-TSC1/2*) (Tapon et al., 2001), (BDRC #80576; RRID:BDSC_80576).

### Luminometry of single larvae

We quantified developmental timing in *D. melanogaster* using a bioluminescence-based assay adapted for individual larvae. To obtain age-matched embryos, young adult flies (1-3 days old) were allowed to acclimate and lay eggs for 2 days at 25 °C in apple agar plates within embryo collection cages. Subsequently, synchronized 0–3 h embryos were obtained by replacing the apple agar plate with a fresh one and allowing egg laying for 3 h. Then, individual embryos were transferred using an eyelash into the wells of a white 96-well plate containing 200 μL per well of luciferin-containig food (0.5% agar (Labotaq cat. # 46046), 6% glucose (Cosela cat. #24379.363), 3% sucrose (IntronBiotech cat. # 10274), 3.2% yeast extract (Eulabor cat. # 48045), 0.5% powder yeast (Sigma cat. #51475), 0.25% Nipagin, 0.25% propionic acid, 200 μM D-Luciferin (Biothema cat. # BT11100). The three final components were added after boiling the medium, once the temperature had dropped below 60 ºC. After loading all wells, plates were sealed with a gas-permeable membrane (Breathe-EASIER™, Diversified Biotech BERM-2000). The membrane is manually perforated (one hole per well) to reduce gas accumulation during measurement acquisition, which also prevents larvae from transferring between wells. Prepared plates were immediately placed in a luminometry reader (Berthold Centro XS3). Luminescence was recorded for 1 s per well at 5-min intervals until pupariation using the MikroWin plate-reader software. All experiments were conducted inside a temperature-controlled incubator (Panasonic MIR-154).

Luminometry data were analysed essentially as previously described (Olmedo et al., 2015). Briefly, molt timing was inferred from temporal changes in luminescence to determine the duration of each developmental stage. To determine the timing of the molts, raw luminescence values were binarized by using as a threshold the 75% of a moving average calculated using a 10 h sliding window. Bioluminescence signals were visualised using R software (version 4.3.1). Transitions in the binarized signal were subsequently detected to identify molt boundaries: transitions from 1 to 0 marked the onset of molting, whereas transitions from 0 to 1 indicated molt completion. This approach allowed automated, high-resolution determination of instar durations and molt timing across individual larvae.

### Temperature regulation

We previously measured the temperature in the plate using a data logger Thermochron iButton DS1921G (Maxim Integrated) and observed that it was approximately 3.5 °C higher than that set at the incubator, due to the production of heat from the luminometer. For all experiments, we adjusted the temperature in the incubator to compensate this difference. The temperatures shown in the data correspond to the temperature experienced by the larvae.

For the evaluation of the effect of temperature on larval development, the embryos were shifted from the maintenance temperature (25ºC) to the experimental temperature at 0 to 3 hours after egg lay, when they were place in the luminometer.

### Arrhenius analysis

To quantify the temperature dependence of *Drosophila* larval development, we applied an Arrhenius breakpoint analysis with a custom-made *R* script following the stepwise framework previously applied in *C. elegans* (Mata-Cabana et al., 2022). The natural logarithm of developmental time was plotted against the inverse absolute temperature (1000/T in K). A reference slope was calculated from a linear regression on three central temperatures (19, 22 and 25°C). Temperatures were then added sequentially, and the slope was recalculated at each step. The first temperature at which the slope deviated by more than 10% from the reference was defined as the Arrhenius break, representing the upper limit of approximate Arrhenius scaling.

### Developmental timing under genetic perturbation

To test the assay with genetically manipulated animals, we ubiquitously reduced TOR signaling by co-expressing UAS-TSC1/2 using *daughterless*-GAL4 in the luciferase background. Control animals were *daughterless*-GAL4 with *ubiquitous-Luciferase* crossed to *w1118* males. Embryos were collected and loaded into luminometry plates as described above. Luminescence was recorded continuously to monitor developmental timing across all larval stages.

### Summary of experimental replicates and number of animals

Figure 1 and supplementary figure 1 contain measurements of 145 animals from 4 independent experiments. The number of animals for each condition in figure 2 and supplementary figure 2 are 93 (16ºC, 2 independent experiments); 129 (19ºC, 2 independent experiments); 93 (22ºC, 2 independent experiments); 145 animals (25ºC, 4 independent experiments); 104 animals (28ºC, 3 independent experiments); 117 animals (28.5ºC, 2 independent experiments); 101 animals (29ºC, 2 independent experiments); and 41 animals (30ºC, 2 independent experiments). All raw data are available in supplementary files (Supplementary file 1, 2). The number of animals for each condition in figure 3 are 26 animals (control group) and 33 (UAS-TSC1/2 group, 2 independent experiments).

### Statistical analysis

Stage durations and developmental timings were derived from automated luminometry traces, as described above. To assess inter-individual variability, we calculated the coefficient of variation (CV) for each larval stage and for total postembryonic development. The differences in CV between stages were analysed with ANOVA followed by Tukey’s post hoc test. The relationship between the duration of larval stages was assessed using Pearson correlation coefficients. Linear regression was applied to assess temperature scaling, including Arrhenius-type relationships for overall developmental rate and individual stage rates. Slopes of fractional stage durations versus temperature were tested against zero to evaluate proportional scaling.

Genetic perturbation experiments were analyzed by comparing stage durations between experimental (UAS-TSC1/2) and control groups using unpaired two-tailed Student’s *t*-tests. Statistical significance was set at p < 0.05. All analyses were performed in GraphPad Prism 10.

## Author Contributions

PRR and MO conceived and designed the study (Conceptualization). DSC, FCL, PRR and NR performed the experiments (Investigation). DSC, FCL, PRR, and MO analyzed and interpreted the data (Formal analysis, Data curation). DSC and MO wrote the original draft of the manuscript (Writing – Original Draft), and all authors contributed to reviewing and editing the manuscript (Writing – Review & Editing). DSC and MO prepared the figures (Visualization). PRR and MO supervised the project (Supervision). All authors read and approved the final manuscript.

## Acknowledgments

We thank the Centro Andaluz de Biología del Desarrollo (CABD) and Acaimo González Reyes for kindly providing the *Drosophila* fly food. We also thank Juan Garrido Maraver and Salvador Herrera for providing *Drosophila* stocks. Stocks obtained from the Bloomington Drosophila Stock Center (NIH P40OD018537) were used in this study.

## Data availability statement

The authors affirm that all data necessary for confirming the conclusions of the article are present within the article, figures, and supplementary files. The raw data for all figures is presented in Supplementary Files 1 and 2.

## Supplementary Material

**Supplementary Figure 1.**
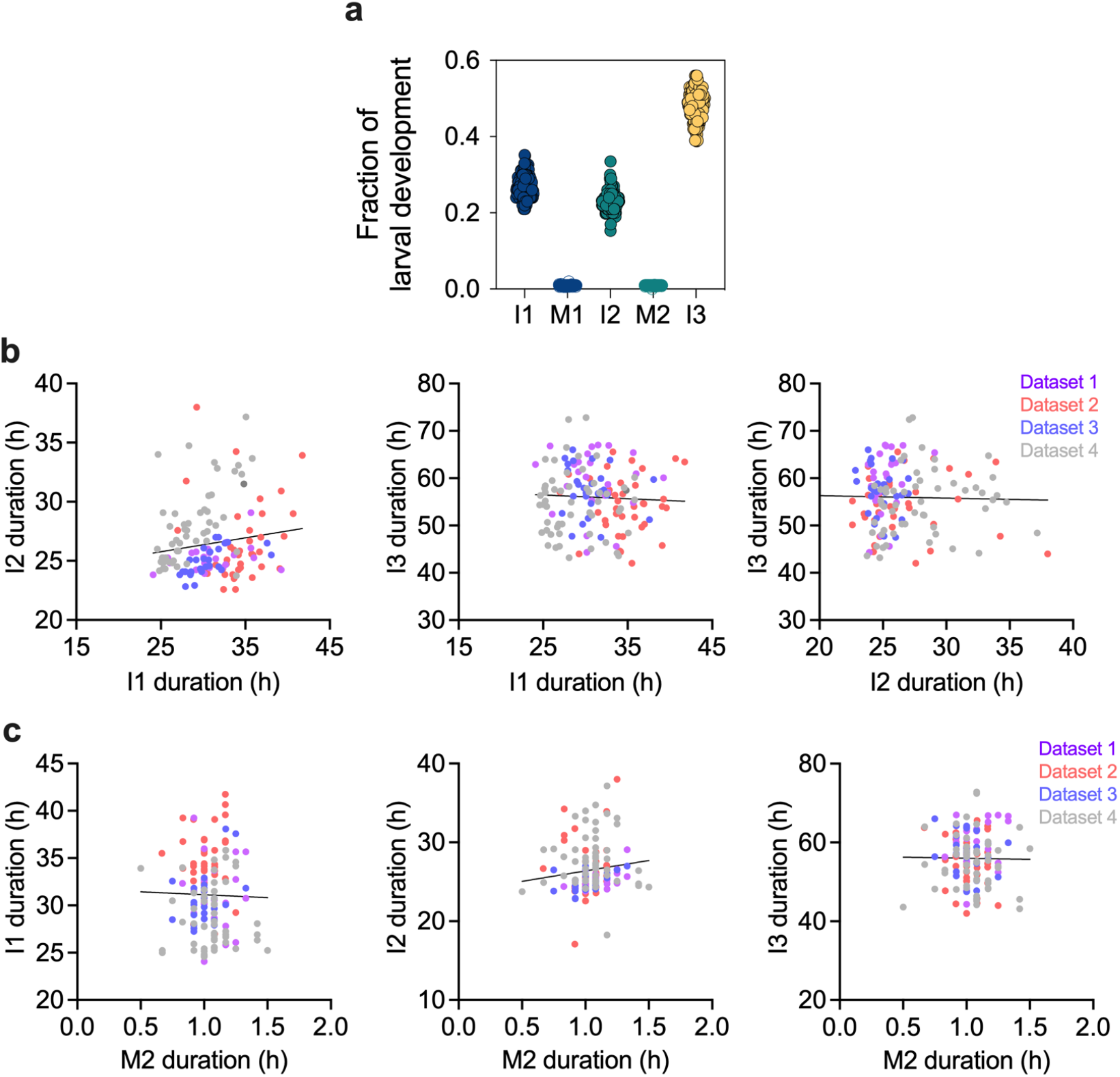
(a) Fraction of total developmental time represented by each stage, showing the proportional contribution of I1, M1, I2, and I3 to overall larval development. (b) Pairwise correlations between the durations of all instars. (c) Pairwise correlations between the durations of M2 vs all instars.

**Supplementary Figure 2.**
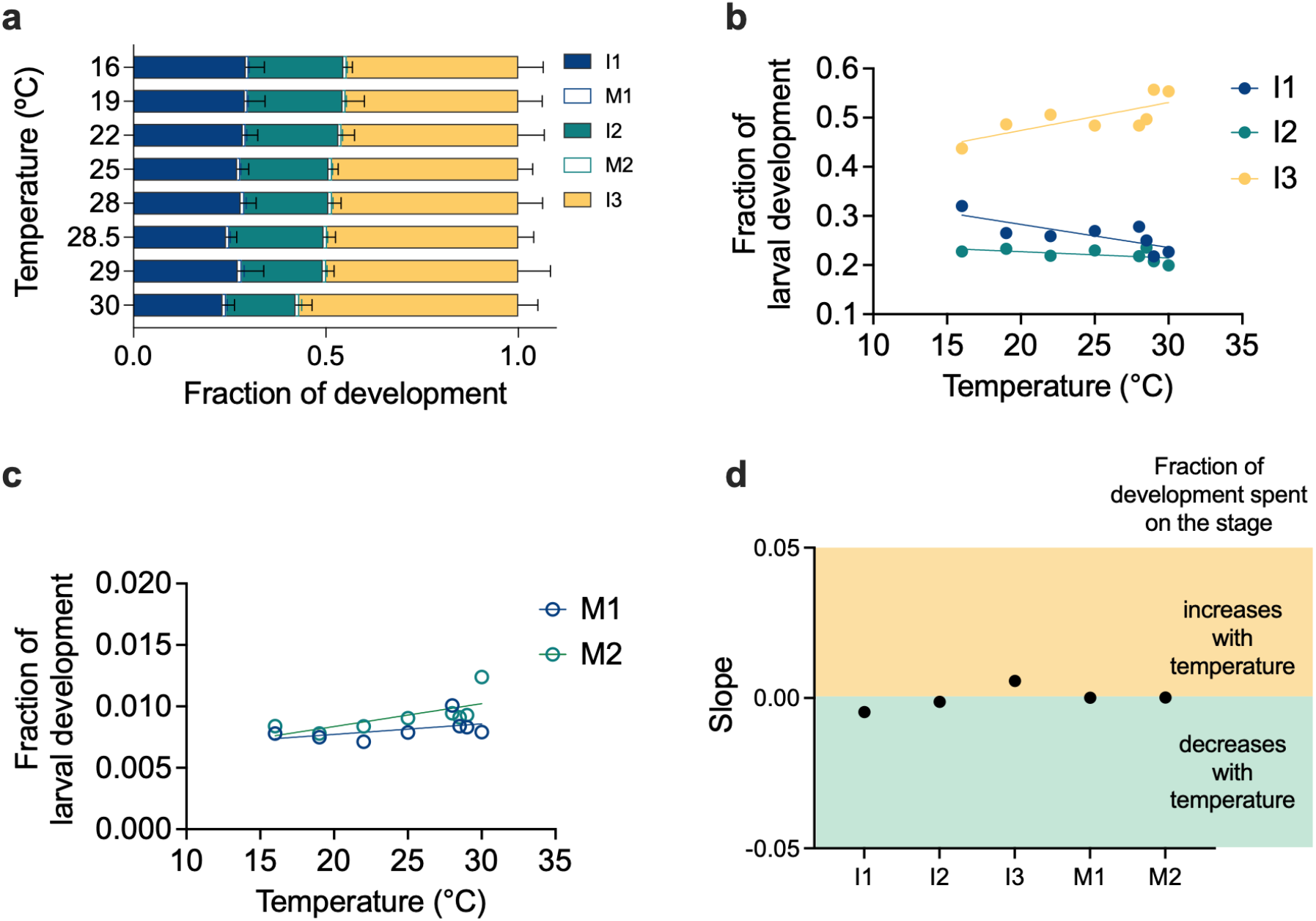
(a) Fraction of total developmental time spent in each stage across temperatures. (b) Datapoints corresponding to panel b for larval instars, highlighting that the relative duration of instars shows no significant temperature-dependent trend. (c) As in panel b, but for molt transitions, demonstrating likewise temperature-invariant fractional timing. (d) Representation of slopes from c and d showing not changes in fraction of development spend on each larva stage (slope values are: −0.0047 for I1; 0.000 for M1; −0.00129 for I2; 0.00019 for M2 and 0.00568 for I3).

Supplementary File 1. Raw data used for figure 1 and 2.

Supplementary File 2. Raw data used for figure 3.

